# Complex cell-state changes revealed by single cell RNA sequencing of 76,149 microglia throughout the mouse lifespan and in the injured brain

**DOI:** 10.1101/406140

**Authors:** Timothy R. Hammond, Connor Dufort, Lasse Dissing-Olesen, Stefanie Giera, Adam Young, Alec Wysoker, Alec J. Walker, Michael Segel, James Nemesh, Arpiar Saunders, Evan Macosko, Robin J. M. Franklin, Xianhua Piao, Steve McCarroll, Beth Stevens

## Abstract

Microglia, the resident immune cells of the brain, rapidly change states in response to their environment, but we lack molecular and functional signatures of different microglial populations. In this study, we analyzed the RNA expression patterns of more than 76,000 individual microglia during development, old age and after brain injury. Analysis uncovered at least nine transcriptionally distinct microglial states, which expressed unique sets of genes and were localized in the brain using specific markers. The greatest microglial heterogeneity was found at young ages; however, several states - including chemokine-enriched inflammatory microglia - persisted throughout the lifespan or increased in the aged brain. Multiple reactive microglial subtypes were also found following demyelinating injury in mice, at least one of which was also found in human MS lesions. These unique microglia signatures can be used to better understand microglia function and to identify and manipulate specific subpopulations in health and disease.

---

Microglia are the resident macrophages of the brain, comprising 10% of brain cells. Not only are microglia active in injury and disease, but they also play critical roles in brain maintenance and development. Microglia, derived from myeloid progenitors in the yolk sac, first arrive in the brain around embryonic day 9 (E9.5) in mice (Ginhoux et al., 2010). Signals in the brain environment shape their maturation by driving broad changes in gene transcription, morphology, and cell number (Butovsky et al., 2014; Gosselin et al., 2014; Lavin et al., 2014; Mass et al., 2016; Matcovitch-Natan et al., 2016).

During this time, microglia also guide neural development, in part by responding to local changes in the brain microenvironment and by interacting with developing neurons. Many of these functional interactions are spatially and temporally controlled and include phagocytosing apoptotic cells, pruning synapses, modulating neurogenesis, and regulating synapse plasticity and myelin formation (Schafer and Stevens, 2015). These distinct functions are often accompanied by regional differences in microglia distribution and morphology, which change as the brain matures. These observations invite the question of whether specialized subpopulations of microglia exist within the brain to carry out these critical, diverse tasks.

In addition to their roles in development, microglia are essential for maintaining the health and function of the brain, as genetic lesions in microglia caused by loss of function mutations in *triggering receptor expressed on myeloid cells 2 (Trem2)* and *colony stimulating factor 1 receptor (Csf1r)* can cause neurodegenerative disease and leukodystrophies, respectively (Paloneva et al., 2000; Rademakers et al., 2011). Moreover, as immune cells, microglia quickly respond to disruptions caused by injury, pathology, or aging (Salter and Stevens, 2017). These responses, often-termed ‘activation’, are defined as any physical or biochemical changes away from the microglial homeostatic state and include rapid proliferation, migration to the site of pathology, phagocytosis of cells and debris, and production of the cytokines and chemokines necessary to stimulate microglia and other brain and immune cells.

Despite the morphological diversity present among microglia in development, health, injury, and disease, microglia have historically been characterized as ‘resting’, ‘M1’ (proinflammatory), or ‘M2’ (anti-inflammatory) - based on simple *in vitro* stimulation methods. Though tools to identifyand manipulate microglia in specific contexts are greatly needed (Ransohoff, 2016), a currently simplistic classification scheme may complicate this by lumping together heterogenous sets of microglia. Identifying and molecularly describing these distinct groups of microglia would help us determine whether microglia assume different profiles based on the type of injury or disease and, in particular, how these states relate to brain development.

Recent high-throughput approaches for single-cell RNA-seq now allow detailed examination of cell state changes and diversity that are reflective of those *in vivo* (Klein et al., 2015; Macosko et al., 2015). A recent study identified a transcriptional signature of microglia surrounding amyloid deposits in a mouse model of Alzheimer’s disease and showed that cells expressing this disease-associated signature (termed disease associated microglia (DAMs)) were present in other mouse models of neurodegeneration and aging (Keren-Shaul et al., 2017).

In this study, we sought to identify the populations of microglia present from embryogenesis to old age and following a local demyelinating injury. We performed high-throughput single-cell transcriptomics of 76,149 mouse microglia and detected many distinct microglial subpopulations with unique molecular signatures that changed over the course of development or in response to injury. We found that microglia were most diverse during early development and became less heterogenous in adulthood, until perturbed by injury or aging. Most of these distinct microglial states were previously unknown and undefined. The detailed molecular signatures identified in our study will lead to a better understanding of the function, signaling, and interaction of distinct microglial subtypes with other brain cells and will facilitate the identification of specific markers that can be used to detect and manipulate microglial states in human health and disease.

## Results

### Isolation of microglia to minimize *ex vivo* activation

In early development, microglia assume a variety of different morphologies and are distributed unevenly in the brain (Karperien et al., 2013). They congregate in specific areas, including the ventricular zone and around growing axon tracts, and not in other areas, like the developing cortex (Squarzoni et al., 2014), suggesting that transcriptionally and functionally different subpopulations of microglia exist. To ask whether microglia are heterogenous during these early time points and to define microglial state changes over time, we isolated whole brains from mice at embryonic day 14.5 (E14.5), early postnatal day 4/5 (P4/P5), the late juvenile stage (P30), adulthood (P100), and old age (18 month) (Fig 1a). To compare healthy development to pathology, microglia were also collected from the white matter of adult mice exposed to a focal demyelinating injury caused by lysolecithin (LPC) injection (Hall, 1972).

**Figure 1.**
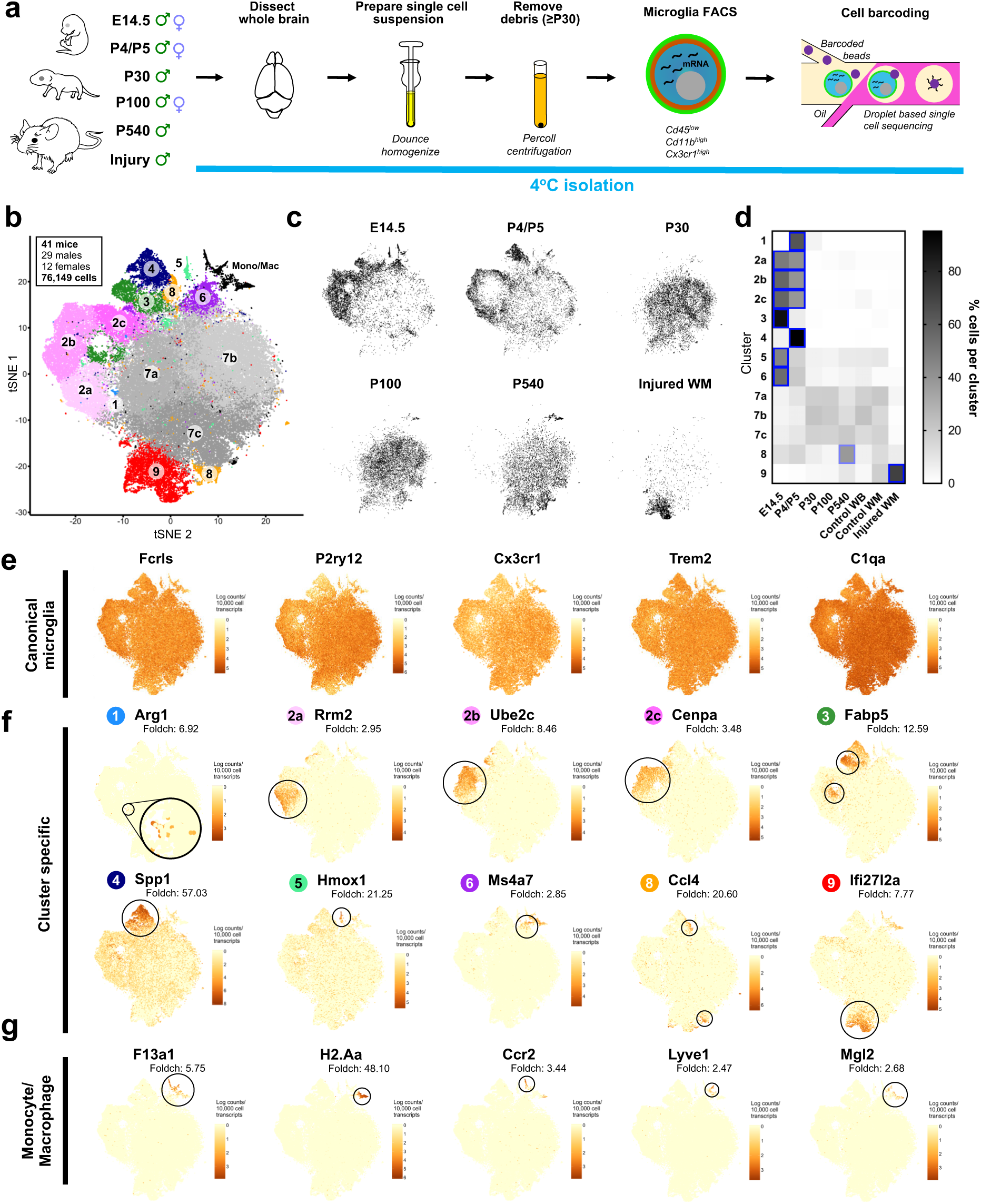
Molecularly distinct subpopulations of microglia peak in number during early development, expand in aging, and emerge following injury. (a) Microglia were isolated from the whole brains of mice from E14.5 until 18 months (P540) and from focal demyelinated white matter lesions (Injury). Microglia were isolated under cold conditions to limit ex vivo activation. Brains were dissociated by dounce homogenization, myelin/debris was removed using a Percoll gradient (on mice older than P5), microglia were purified by FACS (Cd11bhigh, CD45low, Cx3cr1+), and processed using droplet-based single-cell sequencing. (b) tSNE plot of 76,149 cells from 41 animals from all ages and conditions clustered following removal of contaminating cells and batch artifacts. In total, nine microglia clusters and one monocyte/macrophage (Mono/mac)-containing cluster were identified. (c) tSNE plots of cells from each age and condition. For comparison purposes, only male samples were plotted. (d) Heatmap of the normalized percent of cells from each sample assigned to each cluster. Blue squares indicate a significant enrichment (P<0.0001) of cells from a given age in each cluster. Two-way ANOVA with Tukey’s post-hoc analysis. (e) tSNE plot of all 76,149 cells colored for expression (log-transformed UMI counts per 10,000 transcripts) of canonical microglial genes. (f) tSNE plot of expression for genes specifically enriched in each of the microglial clusters. (g) tSNE plot of expression for genes specifically enriched in the monocyte and macrophage cluster.

To minimize *ex vivo* activation and transcriptional activity during the isolation procedure, we generated single-cell suspensions under consistently cold conditions (samples on ice or at 4°C for the duration of the protocol, Fig 1a). Following perfusion, brains were minced and Dounce homogenized; older samples (≥P30) were subjected to Percoll gradient centrifugation to remove debris and myelin. To ensure this step did not affect our recovery of microglial subpopulations, we Percoll purified microglia at P5, when we see considerable microglial diversity, and compared these microglia to those isolated without Percoll - finding no change in the subpopulations present and only a slight shift in the relative percentages of microglia in one of the nine clusters (Supp. Fig 3). Microglia were then FACS-purified by Cd45^low^CD11b^high^Cx3Cr1^high^ expression (Supp. Fig 2) To maximize the fraction of cells that could be recovered and analyzed from these preparations, we utilized a recent commercial implementation of droplet-based RNA-seq analysis (Fig 1a) (Zheng et al., 2017). At least three to four biological replicates per age were collected for a total of 76,149 sequenced microglia from 41 total animals (Fig 1b). Cells were sequenced to comparable sequencing depths (∼40,000-60,000 reads/cell), and after filtering out cells with fewer than 650 transcribed genes, we obtained a similar median unique molecular identifier (UMI) count (∼2100 UMIs/cell) and median gene number (∼1100 genes/cell) in all ages and conditions (Supp. Fig 1a).

To identify transcriptionally distinct microglial subpopulations, we performed dimensionality reduction and clustering using an independent component analysis (ICA)-based approach that has been recently described (Saunders et al., 2018). A first round of clustering was used to identify, curate and remove from analysis contaminating cells (including neurons, endothelial cells, and other cell types) (Supp. Fig 1b,c), which despite FACS purification, constituted a substantial proportion of cells - particularly at younger ages, when neurons were found in our dataset. Non-microglial monocytes and macrophage populations were also found (Cd45^high^ cells) but were left in the clustering analysis to provide a point of reference for comparison with the various microglial states (Fig 1b,g, Mono/Mac cluster). For both rounds of analysis, independent components that captured batch or replicate effects were removed before clustering analysis (Supp. Fig 1d), as previously described (Saunders et al., 2018).

### Distinct microglia subpopulations with unique transcriptional signatures peak in number during early development

Clustering analysis revealed nine unique microglial states across all ages and conditions, including injury (Fig 1b). Cluster sizes ranged from 0.2% of all microglia to as many as 24% of all microglia. Some clusters were dominated by microglia from specific ages, whereas others contained microglia from several ages, indicating that some microglial states are present across development while others are more transitional (Fig 1c,d). We found the greatest microglial diversity at the youngest ages (E14.5 and P5) and considerably less diversity in juveniles (P30) and adults (P100) (Fig 1d). Both aging and injury caused a redistribution of microglial states, including an increase in the percent of cells occupying Clusters 8 and 9, as compared to juveniles and adults (Fig 1d).

Gene expression analysis showed that the canonical microglial genes (*Fcrls, P2ry12, Cx3cr1, Trem2*, and *C1qa*) were highly expressed by most of the analyzed cells, but interestingly, only three (*C1qa, Fcrls, Trem2*) were uniformly expressed in all clusters (Fig 1e), suggesting existing tools and marker definitions need to be updated. Many microglial marker definitions were previously established in adult animals (Butovsky et al., 2014; Hickman et al., 2013), and we found that *P2ry12, Cx3cr1,* and *Tmem119* (not shown) were expressed at much lower levels or not at all in certain clusters of microglia from the developing brain (Clusters 3,4).

In addition to the highly expressed canonical genes, we also identified genes highly enriched in, if not completely unique to, specific microglial states. These unique gene expression patterns show that each microglial state reflects a specific and definable transcriptional program, rather than a simple modulation of commonly expressed microglial genes. Uniquely expressed genes found predominantly at the youngest ages (E14.5 and P4/P5) included arginase 1 (*Arg1,* Cluster 1), ribonucleotide reductase M2 (*Rrm2,* Cluster 2a), ubiquitin-conjugating enzyme E2C (*Ube2c*, Cluster 2b), centromere protein A (*Cenpa*, Cluster 2c), fatty acid binding protein 5, epidermal (*Fabp5,* Cluster 3), osteopontin (*Spp1,* Cluster 4), heme oxygenase 1 (*Hmox1*, Cluster 5), and membrane-spanning 4-domains, subfamily A, member 7 (*Ms4a7,* Cluster 6). Juvenile (P30) and adult (P100) microglia were largely assigned to three central clusters (Cluster 7a-c) that were not defined by any unique genes (Fig 1b). In aged mice, the most enriched subpopulation was defined by the gene chemokine (C-C motif) ligand 4 (*Ccl4,* Cluster 8). Interestingly, Cluster 8 cells were found at most ages, albeit in smaller numbers, with a developmental peak at P5. Cluster 8 formed two spatially distinct groups on the tSNE plot (Figure 1b), which could reflect differences in these cells as the brain ages. Cluster 9 was composed predominantly of microglia from the focal white matter injury and was enriched for the gene interferon, alpha-inducible protein 27 like protein 2A (*Ifi27l2a).* Non-microglial macrophages and monocytes uniquely expressed the genes coagulation factor XIII, A1 subunit (*F13a1,* macrophage), histocompatibility 2, class II antigen A, alpha (*H2-Aa*, macrophage), chemokine (C-C motif) receptor 2 (*Ccr2*, monocyte), lymphatic vessel endothelial hyaluronan receptor 1 (*Lyve1,* macrophage), and macrophage galactose N-acetyl-galactosamine specific lectin 2 (*Mgl2,* macrophage), genes that were barely expressed, if at all, by microglia (Fig 1g).

Together, these data demonstrate that microglia exist in multiple definable states that change over the course of development, aging, and injury. Although further analysis will be needed to establish whether these states are transient or represent the terminal differentiation of different subsets of microglia, these data provide a comprehensive map of these changes and identify genes that define several of these specific microglial states.

### A novel population of embryonic Ms4a-expressing microglia share a similar transcriptional profile with brain border macrophages

Microglia and brain border macrophages (which reside in the perivascular space, meninges, and choroid plexus) are derived from the same pool of yolk sac hematopoietic progenitors and migrate to the brain at the same time in development (Goldmann et al., 2016). It is not until microglia infiltrate the brain parenchyma and are exposed to brain-derived signals that they achieve their unique identity. However, it is still unclear when microglia diverge from their brain border neighbors and how quickly they differentiate. At E14.5, we identified a distinct subpopulation of microglia that uniquely express *Ms4a7* and share a similar transcriptional profile with brain border macrophages (Fig 2, Cluster 6). Cluster 6 microglia were enriched for canonical markers of brain border macrophages, including mannose receptor, C type 1 (*Mrc1*), but lacked other markers like *F13a1* and *Lyve1* (Fig 1g). It is possible that Cluster 6 microglia downregulate these genes as they enter the brain, but this question will need closer investigation.

**Figure 2.**
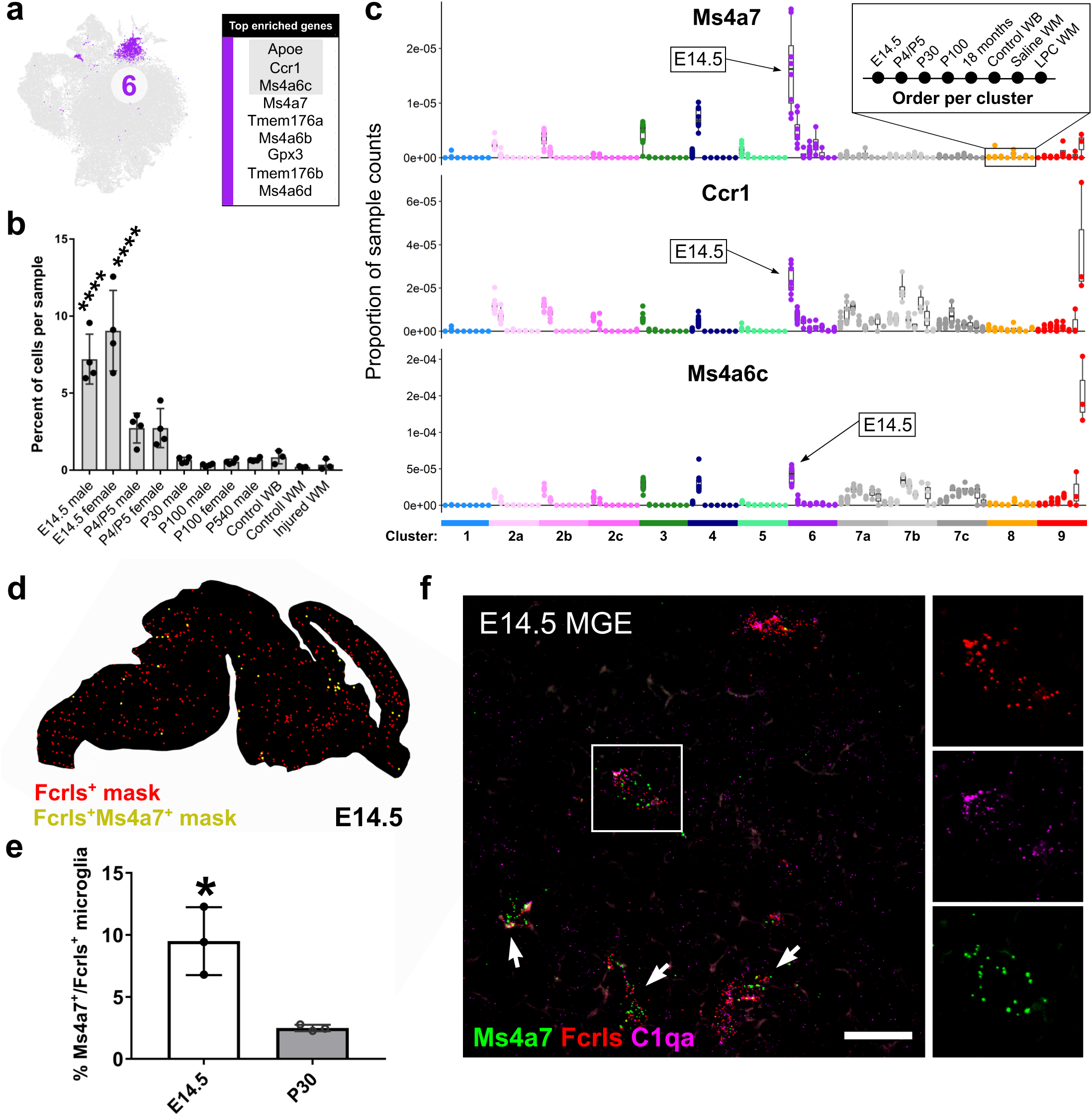
Identification of Ms4a7-expressing microglia in the embryonic brain that resemble brain border macrophages. (a) tSNE plot of Cluster 6 microglia and a table of the top nine enriched genes in that cluster. Gray outlined genes are plotted in (c). (b) Plot of the percent of cells per sample that were assigned to Cluster 6. ****P<0.0001. ANOVA with Tukey’s post-hoc analysis. (c) Plot of the proportion of normalized UMI counts per sample (summed cell counts) for cells assigned to each cluster for the top genes in Cluster 6. Sample order per cluster is listed in the inset. Sample/ages that are enriched for a given gene are denoted by an arrow. (d) Representive image of masked microglia identified by our high-throughput cell quantification pipeline in a E14.5 saggital brain section. Cells were marked as single positive (Fcrls+ only, red) or double positive (Fcrls+Ms4a7+, yellow). (e) Quantification of the percent of Fcrls+ microglia that also expressed Ms4a7 using high-throughput automated counting of smFISH probe cells. 3-4 saggital brain sections per animal in 3 animals were quantified at E14.5 and P30. *P<0.05, Unpaired t-test. (f) High-magnification confocal image of E14.5 brain section of the intermediate zone of the telencephalon stained by smFISH with pan-microglia probes Fcrls and C1qa and Cluster 6-specific probe Ms4a7. Scale bar = 20 microns.

In addition to *Ms4a7*, Cluster 6 microglia were enriched for other Ms4a family members including *Ms4a6c, Ms4a6b, Ms4a6d, Tmem176a,* and *Tmem176b* (Fig 2a). The Ms4a family genes are transmembrane chemosensors (Greer et al., 2016), some of which regulate immune cell function (Eon Kuek et al., 2016). However, the function of the Ms4a family members in cell- and tissue-specific contexts is largely unresolved. Interestingly, Ms4a family members are associated with Alzheimer’s risk (Hollingworth et al., 2011; Ma et al., 2015; Naj et al., 2011), but their role in the disease is not understood. Cluster 6 microglia were also enriched for chemokine (C-C motif) receptor 1 (*Ccr1*), which regulates immune cell migration and functional states (Chou et al., 2010; Furuichi et al., 2008; Mahad et al., 2004). This finding might indicate *Ccr1* is an important receptor for microglial migration or function in the developing brain.

Cluster 6 microglia were highly enriched at E14.5 versus any other age or condition analyzed, as approximately 10% of all E14.5 microglia (but no more than 3% of microglia at any other age) fall into Cluster 6 (Fig 2b). In support of an early embryonic enrichment of these cells, *Ms4a7* gene expression in Cluster 6 was most concentrated at E14.5 (Fig 2c). To ensure Cluster 6 microglia were localized in the brain parenchyma, we performed single-molecule fluorescent *in situ* hybridization (smFISH) followed by an automated high-throughput quantification (see methods) of *Ms4a7* expression in microglia in the E14.5 and P30 brains. Microglia were marked by *Fcrls*, which was uniformly expressed by all microglia in every cluster (Fig 2d). *Ms4a7*^+^ microglia were sparsely distributed throughout the embryonic brain (Fig 2d,f), with a population of *Ms4a7*^+^ macrophages at the brain borders (not shown). *Ms4a7*^+^ microglia were strongly enriched at E14.5 (Fig 2e), and the small percentage of *Ms4a7*^+^ cells at P30 likely represent perivascular macrophages. These differences found by smFISH closely mimic the percentages of *Ms4a7*^+^ microglia found in our sequencing data and confirm that these cells are present in the developing brain.

Altogether, these results identify Cluster 6 microglia as a novel subpopulation that is enriched in the embryonic brain parenchyma and bears considerable transcriptional overlap with brain border macrophages. These cells might give rise to perivascular macrophages or could represent a precursor to mature microglia, but lineage tracing will be necessary to tease those possibilities apart.

### Specialized axon tract-associated microglia appear during a restricted developmental window

In the early postnatal brain, microglia regulate the growth and fasciculation of axons and can refine synapses in a circuit- and region-specific manner (Schafer and Stevens, 2015; Squarzoni et al., 2014). At P4/P5, we found one major microglial state (Cluster 4) that was barely found at any other time point (Fig 3a,b,c). These microglia were highly enriched for the gene secreted phosphoprotein 1 (*Spp1*), also known as osteopontin (Fig 3a,c). Other enriched genes included insulin-like growth factor 1 (*Igf1*), glycoprotein (transmembrane) NMB (*Gpnmb*), and immunomodulators from the galectin family, galectin-1 (*Lgals1*) and galectin-3 (*Lgals3).* These cells, which assume an amoeboid morphology (data not shown), also upregulate the lysosomal markers lysosomal-associated membrane protein 1 (*Lamp1*) and *Cd68*, the latter of which is increased in phagocytic microglia (Schafer and Stevens, 2015). Interestingly, smFISH for *Spp1* showed that these cells only resided in the subcortical axon tracts of the forebrain, as well as in distinct clusters in the axon tracts of the cerebellum (Fig 3d-f). The axon tracts where Cluster 4 microglia were concentrated will eventually become heavily myelinated, but Cluster 4 microglia are largely gone before myelination occurs. Interestingly, there was striking diversity of expression on a cell-to-cell basis in these areas, with some microglia expressing high levels of *Spp1* adjacent to neighbors that had no detectable *Spp1* expression (Fig 3f). These data suggest microglial heterogeneity also exists on a very local scale.

**Figure 3.**
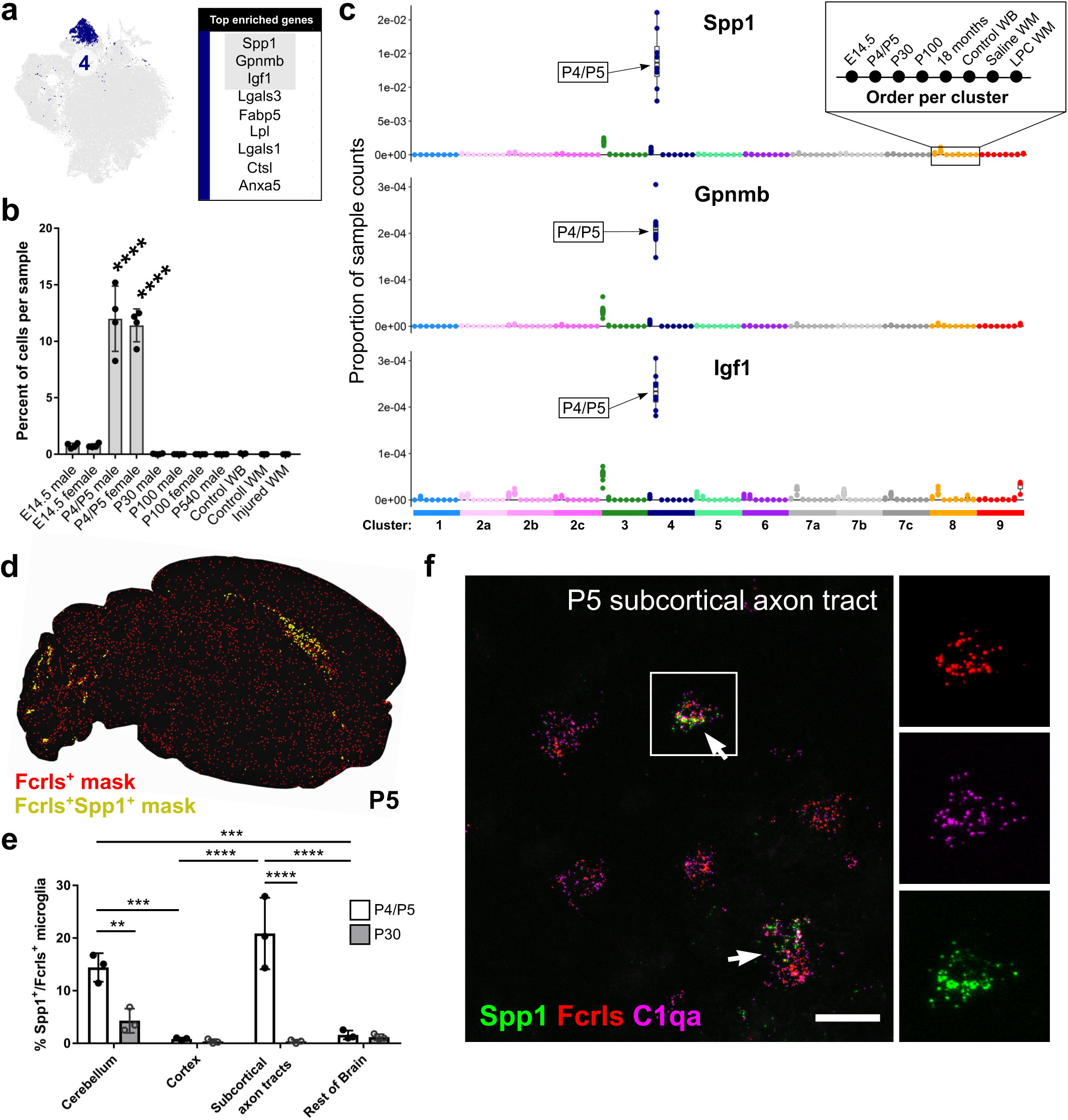
Spp1-expressing microglia densely occupy premyelinated axon tracts in the early postnatal brain. (a) tSNE plot of Cluster 4 microglia and a table of the top nine enriched genes in that cluster. Gray outlined genes are plotted in (c). (b) Plot of the percent of cells per sample that were assigned to Cluster 4. ****P<0.0001. ANOVA with Tukey’s post-hoc analysis. (c) Plot of the proportion of normalized UMI counts per sample (summed cell counts) for cells assigned to each cluster for the top genes in Cluster 4. Sample order per cluster is listed in the inset. Sample/ages that are enriched for a given gene are denoted by an arrow. (d) Representive image of masked microglia identified by our high-throughput cell quantification pipeline in a P5 saggital brain section. Cells were marked as single positive (Fcrls+ only, red) or double positive (Fcrls+Spp1+, yellow). (e) Quantification of the percent of Fcrls+ microglia that also expressed Spp1 using high-throughput automated counting of smFISH probe cells. 3-4 saggital brain sections per animal in 3 animals were quantified in four brain regions at P5 and P30. ****P<0.0001, ***P<0.001, Two-way ANOVA, Tukey’s post-hoc analysis. (f) High-magnification confocal image of the P5 subcortical axon tracts in the forebrain stained by smFISH with pan-microglia probes Fcrls and C1qa and Cluster 4-specific probe Spp1. Scale bar = 20 microns.

In conclusion, this Spp1+ microglia population are highly concentrated specifically in the axon tracts of the premyelinated brain. Their amoeboid morphology and their enrichment of genes associated with immune cell activation, lysosomal activity, and phagocytosis suggest that Cluster 4 microglia might engulf material in these regions. Further experiments will be needed to explore what this material could be, but their restricted tempo-spatial appearance suggests that their activity is highly regulated.

### Microglia expansion and distribution fueled by metabolically active and proliferative microglia in early development

Microglia achieve their final cell numbers and distribution in postnatal development after brain development is well underway (Squarzoni et al., 2014). At early embryonic ages, microglia are far less abundant than in adulthood, when they constitute 10% of all brain cells. Rapid expansion of microglia also occurs in response to injury or disease, and their self-renewal capacity is sufficient to replenish the population within days following depletion of all but 1-2% of microglia (Elmore et al., 2014). Interestingly, microglia are one of the few cell types in the brain that do not have a known progenitor pool dedicated to developmental expansion or renewal. Here, we uncovered several large microglial subpopulations in the embryonic and early postnatal brain that expressed markers associated with metabolic activity, cell proliferation, cell growth, and cell motility that could underlie the brain colonization process (Supp. Fig 5).

The first subpopulation of microglia belong to Cluster 3 and uniquely express the gene *Fabp5*. Cluster 3 microglia were highly enriched for several other genes including macrophage migration inhibitory factor (*Mif*), lactate dehydrogenase A (*Ldha*), and triosephosphate isomerase 1 (*Tpi1*) (Fig 4a,c). *Mif* and *Fabp5* have both been linked to cell growth, motility, inflammation, and immunomodulation in macrophages and other cells (Calandra and Roger, 2003; Kannan-Thulasiraman et al., 2010; Liu et al., 2010). Other enriched transcripts in Cluster 3 microglia were associated with glycolysis, suggesting an altered metabolic profile in these cells (Supp. Fig 5), a feature that is consistent with other cell types in their early, less-differentiated states. Interestingly, a shift to glycolysis from oxidative phosphorylation is a key metabolic alteration in inflammatory macrophages (Mills et al., 2016). Cluster 3 microglia were found almost exclusively at E14.5, although a small percentage were also present at P4/P5 (Fig 4b). At E14.5, approximately 10-15% of all microglia belonged to Cluster 3 (Fig 4b). Surprisingly, the top genes, including *Fabp5, Mif, Ldha,* and *Tpi1,* in Cluster 3 microglia were also partially enriched in the Cluster 4 *Spp1*^+^ microglia found at P4/P5, albeit to a lower extent (Fig 4c). Conversely, Cluster 3 microglia were partially enriched for Cluster 4 markers *Spp1, Gpnmb,* and *Igf1* (Fig 3c), suggesting an overlapping transcriptional signature and possible relationship between the two states.

**Figure 4.**
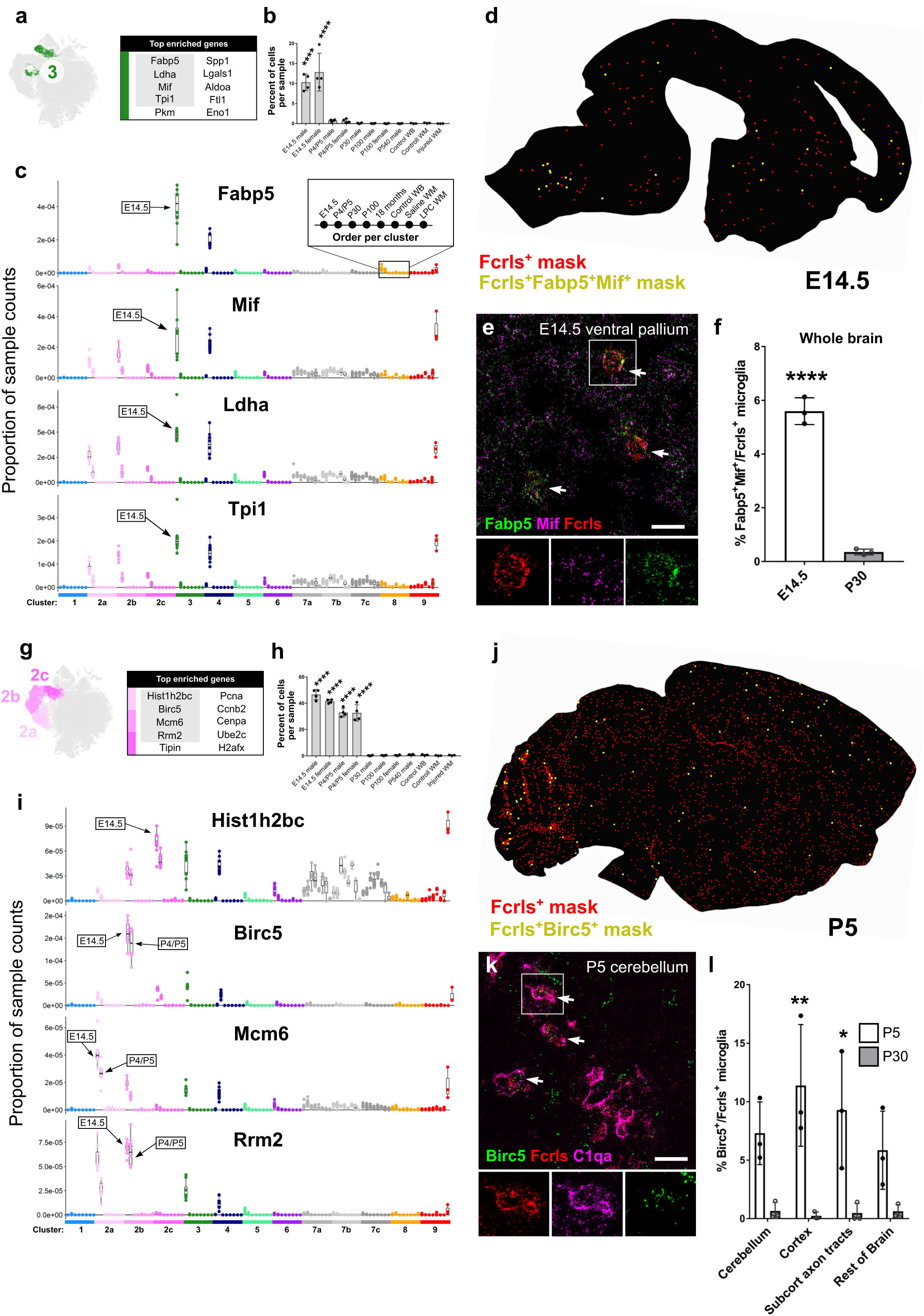
Metabolically active and proliferative microglial subpopulations dominate early brain development. (a) tSNE plot of Cluster 3 microglia and a table of the top nine enriched genes in that cluster. Gray outlined genes are plotted in (c). (b) Plot of the percent of cells per sample that were assigned to Cluster 3. ****P<0.0001. ANOVA with Tukey’s post-hoc analysis. (c) Plot of the proportion of normalized UMI counts per sample (summed cell counts) for cells assigned to each cluster for the top genes in Cluster 3. Sample order per cluster is listed in the inset. Sample/ages that are enriched for a given gene are denoted by an arrow. (d) Representive image of masked microglia identified by our high-throughput cell quantification pipeline in a E14.5 saggital brain section. Cells were marked as single positive (Fcrls+ only, red) or double positive (Fcrls+Fabp5+Mif+, yellow). (e) High-magnification confocal image of the P5 subcortical axon tracts in the forebrain stained by smFISH with pan-microglia probes Fcrls and C1qa and Cluster 3-specific probes Fabp5 and Mif. Scale bar = 20 microns. (f) Quantification of the percent of Fcrls+ microglia that also expressed Fabp5 and Mif by high-throughput automated counting of smFISH probe cells. 3-4 saggital brain sections per a<u>nim</u>al in 3 a<u>nim</u>als were quantified at E14.5 and P30. ****P<0.0001, ***P<0.001, Two-way ANOVA, Tukey’s post-hoc analysis. (g) tSNE plot of Cluster 2 (a-c) microglia and a table of the top ten enriched genes in those clusters. Gray outlined genes are plotted in (c). (h) Plot of the percent of cells per sample that were assigned to Cluster 2. ****P<0.0001. ANOVA with Tukey’s post-hoc analysis. (i) Plot of the proportion of normalized UMI counts per sample (summed cell counts) for cells assigned to each cluster for the top genes in Cluster 2. Sample order per cluster is listed in the inset. Sample/ages that are enriched for a given gene are denoted by an arrow. (j) Representive image of masked microglia identified by our high-throughput cell quantification pipeline in a P5 saggital brain section. Cells were marked as single positive (Fcrls+ only, red) or double positive (Fcrls+Birc5+, yellow). (k) High-magnification confocal image of the P5 subcortical axon tracts in the forebrain stained by smFISH with pan-microglia probes Fcrls and C1qa and Cluster 2b-specific probe Birc5. Scale bar = 20 microns. (l) Quantification of the percent of Fcrls+ microglia that also expressed Birc5 by high-throughput automated counting of smFISH probe cells. 3-4 saggital brain sections per animal in 3 animals were quantified in four brain regions at P5 and P30. ****P<0.0001, ***P<0.001, Two-way ANOVA, Tukey’s post-hoc analysis.

*Mif* and *Fabp5* were both broadly expressed in the E14.5 brain, so both were used as smFISH probes to increase the confidence of colocalization with the microglial marker *Fcrls. Fabp5+Mif*^+^ microglia were distributed throughout the E14.5 brain and often formed small clusters (Fig 4d). smFISH quantification confirmed the striking decrease in the number of *Fabp5+Mif* microglia between E14.5 and P30 (6% vs. <1% of all microglia) (Fig 4e,f). Together, these data show that this unique population of *Fabp5+Mif*^+^ microglia was found almost exclusively in the embryonic brain and were not restricted to a particular brain region.

At E14.5, the most populous microglial states were proliferative (Clusters 2a,2b,2c) and comprised a total of approximately 40% of all E14.5 microglia and 35% of P5 microglia (Fig 4g,h). We grouped these clusters together because of their transcriptional similarity and because it is likely that each of the proliferative clusters represent microglia at different stages of the cell cycle. Indeed, pathway enrichment for M-phase, S-phase, and G-phase were represented to differing degrees in each subgroup (Supp. Fig 5). The top cluster markers for Cluster 2a were minichromosome maintenance complex component 6 (*Mcm6*), proliferating nuclear cell antigen (*Pcna*), and *Rrm2.* For Cluster 2b, they were *Ube2c*, baculoviral IAP repeat-containing 5 (*Birc5*), and H2A histone family, member X (*H2afx*), and for Cluster 2c they were histone cluster 1, H2bc (*Hist1h2bc*), cyclin B2 (*Ccnb2*), and *Cenpa.* Markers for each of the proliferative populations were comparably enriched at E14.5 and P5 and were highly specific to the proliferative clusters (Fig 4i). By age P30 and onward, only a few percent of cells fell into these proliferative clusters (Fig 4h), suggesting that microglia divide at extremely low levels at older ages, an assertion supported by recent publications (Fuger et al., 2017).

Interestingly, the top cluster markers in Clusters 2 and 3 also shared some transcriptional overlap. For example, *Mif, Ldha,* and *Tpi1* were somewhat enriched in the E14.5 and P5 proliferative microglia (Fig 4i). The same is true for the cell proliferation genes, which were enriched to a smaller extent in the Cluster 3 *Fabp5+Mif*^+^ microglia (Fig 1e,4c). Interestingly, a portion of the Cluster 3 microglia physically cluster with the proliferative microglia (Fig 1b) and were also enriched for degradation pathways for mediators of proliferation (Supp. Fig 5), which could indicate that these cells flexibly enter and exit a proliferative state.

Together, these two subpopulations of embryonic and early postnatal microglia likely play essential roles in controlling how microglia populate the brain. The widespread distribution of both proliferative and metabolically active microglia suggests that both cell subtypes give rise to mature microglia in the adult brain, but lineage tracing studies will be needed to track their progression

### Sex has no impact on microglial diversity or the number of cells in each subpopulation

Recent evidence has uncovered sex differences in microglial gene expression and function in the normal and challenged developing and adult brain (Hanamsagar et al., 2017; Sorge et al., 2015; Thion et al., 2018). To determine whether sex had any effect on microglial diversity, we compared microglia from male and female mice at three major developmental ages: E14.5, P4/P5, and P100 (Supp. Fig 4). A total of 49,445 cells were clustered together across all the ages and sexes. We were able to recognize all of the same clusters that were identified in the larger analysis in Figure 1, and the clusters were colored the same way for comparison purposes (Supp. Fig 4a). As expected, microglia from males uniquely expressed the Y chromosome gene *Eif2s3y* and microglia from females expressed the X inactivation gene *Xist* (Supp. Fig 4f). We found almost no difference in the clustering between the male and female samples (Supp. Fig 4b-d), as measured by the number of cells per sample in each cluster (Supp. Fig 4c) and the normalized number of cells occupying each cluster (Supp. Fig 4d).

The only observed difference between the sexes was in Cluster 6, which was enriched in female samples. Cluster 6 is the smallest identified microglial cluster (∼0.5% of microglia), was observed only at P4/P5 (Supp. Fig 4c-d, Fig 1d), and was enriched for the genes *Cd74*, chemokine (C-C motif) ligand 24 (*Ccl24*), and *Arg1.* Of note, *Arg1* is a common marker of peripheral ‘M2’ macrophages (Murray et al., 2014), but we show in our dataset that microglia only express *Arg1 in vivo* in a very small subset of cells. Further analysis will be needed to validate and better understand this extremely small population of microglia and why the number of these microglia would be higher in females than in males. Altogether, microglial diversity was largely unaffected by sex during normal development. Given the findings of sex differences in models of injury, pathology, and germ-free conditions, it is possible that sexual dimorphism in microglia emerge under conditions of challenge (Lenz and McCarthy, 2015; Thion et al., 2018).

### Small populations of inflammatory and interferon-responsive microglia emerge in the aged brain

As the brain ages, it becomes an increasingly challenging environment, characterized by the accumulation of oxygen free radicals, compromise of blood-brain barrier integrity, and loss of functional synapses (Montagne et al., 2015; Murman, 2015; Poon et al., 2004). Microglia may exacerbate some of these deleterious processes by driving or perpetuating brain inflammation through increased expression of inflammatory molecules (Gabuzda and Yankner, 2013; Grabert et al., 2016; Hickman et al., 2013). However, the extent of the microglial response to aging has not been comprehensively studied, and it remains unclear how, where (in the brain), and to what extent microglia change with normal aging. Surprisingly, aging did not lead to the appearance or disappearance of any clusters, but rather caused a progressive expansion of clusters that are typically present at low levels in adolescent and adult samples (P30 and P100, Fig 1d). To more deeply assess how aging impacts microglial states, we performed a direct clustering comparison between microglia from adult (P100) and 18-month-old (P540) mice (Fig 5a). Four microglia clusters were identified as being enriched in aging mice (Aging Clusters 1a, 1b, 2, and 4), along with one brain border macrophage cluster (Aging Cluster 3), which expressed the macrophage-specific markers *F13a1* and *Mgl2* (Fig 1f). Aging Clusters 2, 3 and 4 were enriched 2-4 times in the P540 samples versus P100 controls (Fig 5b,c)

Aging Cluster 2 cells were enriched for a number of inflammatory signals, which were not normally expressed by other populations of microglia *in vivo* (Fig 5d, Aged Cluster 2). They were enriched for *Lgals3,* cystatin F (*Cst7*), chemokines *Ccl4* and *Ccl3,* as well as the inflammatory cytokine interleukin 1 beta (*Il1b*) (Fig 5d). *Ccl4,* also known as macrophage inflammatory protein-1β (Mip-1β), is the ligand for chemokine receptor type 5 (*Ccr5*), which regulates the trafficking and effector functions of diverse populations of immune cells (Weber et al., 2001). Interestingly, *Ccl4*^+^ microglia were present at very low levels in the healthy developing brain, with a small developmental peak at P5 before spiking to their highest level in the P540 brain (Fig 1d). This increase, coupled with the overall increase in the inflammatory environment in the aged brain (Gabuzda and Yankner, 2013), suggests that this small subpopulation of microglia contributes to age-related brain inflammation; however manipulation of these microglia will be essential for testing this hypothesis.

**Figure 5.**
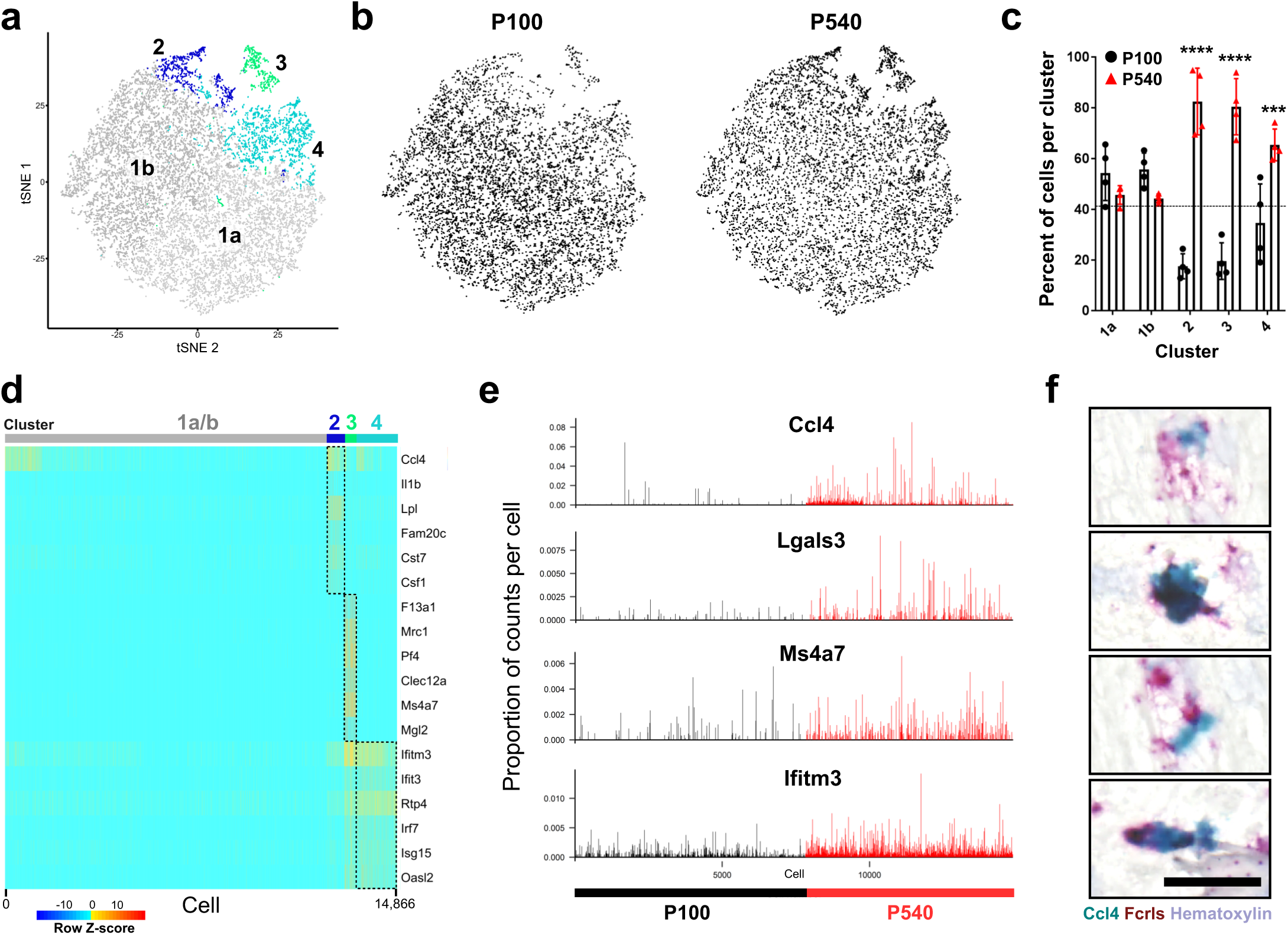
Aging drives an immunogenic state in microglia subpopulations. (a) tSNE plot of 14,866 microglia from P100 and P540 (18 month) male mice (n = 4 per age) shows four microglia clusters (1a, 1b, 2, 4) and one cluster of brain border macrophages (3). (b) Locations of cells from each age on the same tSNE plot as in (a). (c) Plot of the normalized percent of cells from each cluster derived from each age. ****P<0.0001, ***P<0.001, Two-way ANOVA, Tukey’s post-hoc analysis. (d) Heatmap of gene expression in each of 14,886 cells in each of the clusters from (a). Plotted genes are some of the top genes enriched for each cluster. Genes expression scaled by row for plotting and comparison purposes. Z-score represents the number of standard deviations from the mean following scaling. (e) Plot of the number of log-transformed UMI counts per 10,000 cell transcripts in all 14,886 microglia isolated from P100 and P540 samples. (f) Representative high-magnification image of Fcrls+ (red) and Fcrls+Ccl4+ (red and turquoise) cells in the P540 hindbrain. Scale bar = 25 microns.

To localize these Aged Cluster 2 microglia, we performed smFISH for *Ccl4* and *Fcrls* on P100 and P540 brain sections using a non-fluorescent chromogenic method to avoid autofluorescence from lipofuscin granules found in the lysosomes of aged microglia (Xu et al., 2008). We found evidence of *Fcrls+Ccl4*^+^ cells throughout the adult and aged brain (Fig 5f).

Aging Cluster 4 was enriched for interferon-response genes including interferon induced transmembrane protein 3 (*Ifitm3*), receptor transporter protein 4 (*Rtp4*), and 2’-5’ oligoadenylate synthetase-like 2 (*Oasl2*) (Fig 5d,e). Age-related activation of interferon-response genes has been previously reported in the choroid plexus (Baruch et al., 2014) and in microglia (Grabert et al., 2016), but our findings indicate that this profile is actually restricted to a small subset of microglia (Fig 5a,c). Interferon-response genes can modulate inflammation (Baruch et al., 2014), so it is plausible that Aging Cluster 4 microglia also play a role in contributing to the inflammatory tone of the aged brain. Importantly, interferon is produced in response to damage-associated signals such as HMGB1 and cell-free nucleic acids. Therefore, the detection of localized populations of microglia expressing interferon-response genes may assist in identifying foci of neuronal injury or degeneration (Mathys et al., 2017).

Altogether, our data show that aging triggers a shift towards a more immunogenic profile including an increase in inflammatory microglia and interferon-responsive microglia. Even with the shift, the number of microglia that occupy these states form only a small fraction of microglia, suggesting that the vast majority of microglia are unaltered by aging and that local cues could drive state changes rather than a brain-wide environmental shift. The widespread, but sparse, distribution of the *Ccl4*^+^ cells in the aged brain could mean that highly localized events like blood brain barrier compromise (Montagne et al., 2015) or microinfarcts (Smith et al., 2012) trigger these responses.

### Diverse microglial activation responses are triggered in demyelinated mouse lesions and human MS tissue

Research has often referred to microglia that respond to injury or pathology as ‘activated’, a catch-all term for biochemical and physical divergence from a homeostatic state. Activated microglia have been observed in almost every neurological disease including both neurodevelopmental and neurodegenerative disorders (Salter and Stevens, 2017). However, it is still unclear whether or how microglia tailor their response to specific injury types or whether distinct populations of microglia exist in pathological tissue, as we lack both the markers and tools to identify and characterize different activation states, should they exist.

To begin addressing these questions, we used an injury model in which focal demyelination of the subcortical white matter in mice is triggered by injection of lysolecithin (LPC). LPC-induced demyelination is frequently used to study myelin repair and recapitulates many aspects of the demyelinated lesions found in patients with multiple sclerosis (MS) (Hammond et al., 2014). We used this model for several reasons: first, the time course of injury and repair are well established and controlled; second, microglia play a dynamic role in physiological and injury-induced remyelination (Miron et al., 2013); third, microglia respond robustly and rapidly to demyelination in this model; and finally, microglial activation is largely confined to the lesion where the demyelination has occurred. We focused our analysis on 7 days post-lesion (7 dpl), a time point at which microglia are highly responsive and the lesion is undergoing a transition from myelin debris cleanup to the initial phases of remyelination (Fig 6a).

**Figure 6.**
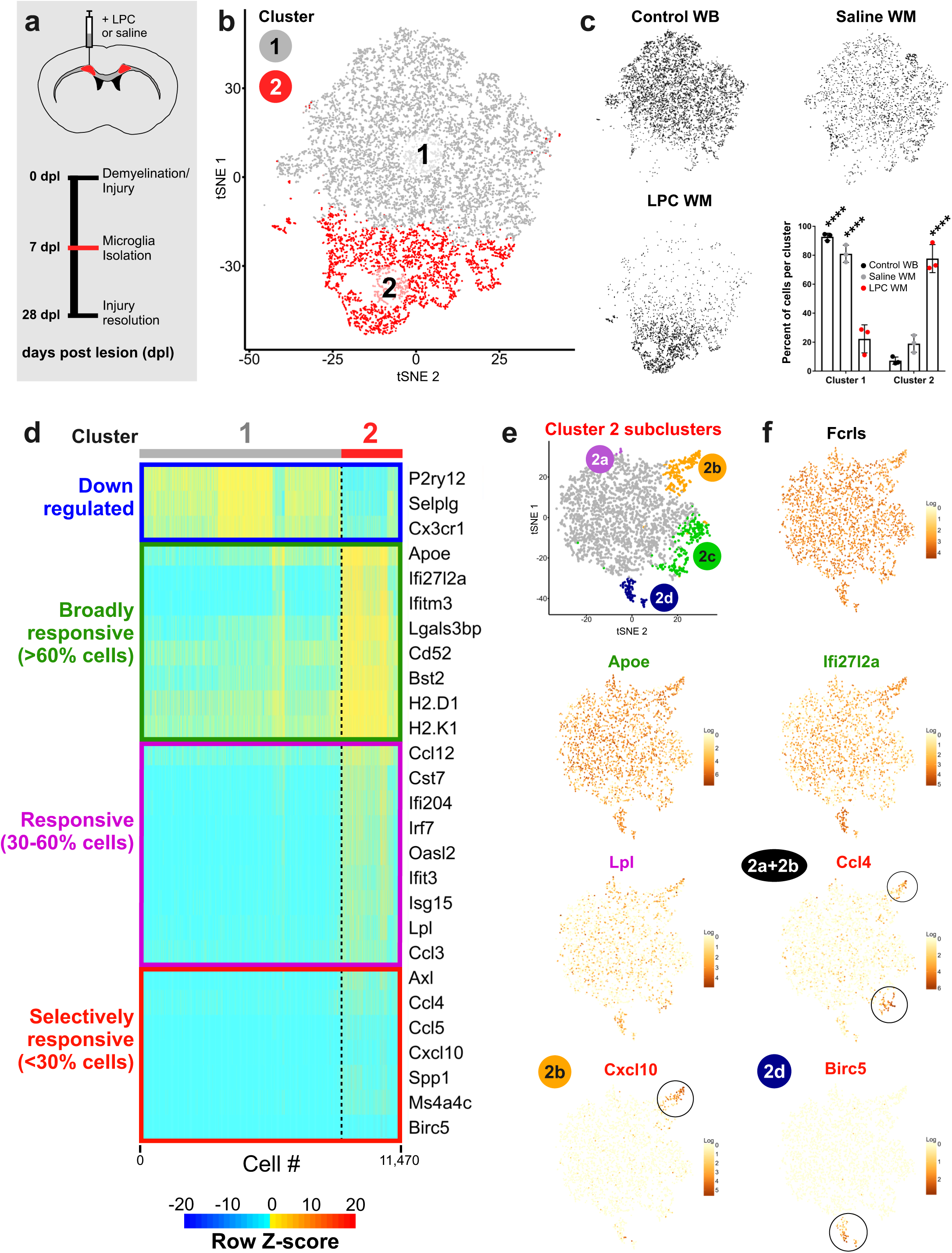
Injury-responsive microglia in demyelinated lesions exhibit multiple activation states. (a) The subcortical white matter of adult male mice was injected with the demyelinating agent lysolecithin (LPC) or saline and microdissected after 7 days post lesion/injection (dpl). Microglia from the demyelinated or control white matter were then isolated and sequenced using the protocols defined in Figure 1a. (b) tSNE plot of 11,470 microglia from LPC-injected white matter, saline-injected white matter, or whole-brain adult (P100) samples reveals two microglia clusters. N = 3 mice per condition. (c) tSNE plots of cells from each condition and a plot of the normalized percent of cells from each sample assigned to each cluster. ****P<0.0001, Two-way ANOVA, Tukey’s post-hoc analysis. (d) Heatmap of gene expression in each of 11,470 cells in each of the clusters from (b). Plotted genes were grouped by genes that were downregulated in Cluster 2 or the percent of cells that upregulated a gene in Cluster 2 - including broadly responsive genes that were upregulated in over 60% of Cluster 2 cells, responsive genes that were upregulated in 30-60% of Cluster 2 cells, and selectively responsive genes that were upregulated in less than 30% of Cluster 2 cells. Genes expression scaled by row for plotting and comparison purposes. Z-score represents the number of standard deviations from the mean following scaling. (e) Cells from Cluster 2 were subclustered to identify different subpopulations of activated microglia. tSNE plot shows four subclusters of Cluster 2 microglia (2a-2d). (f) tSNE plots of subclustered cells colored for expression (log-transformed counts per 10,000 transcripts) of genes that were broadly expressed or upregulated in the majority of Cluster 2 microglia (Fcrls, Apoe, Ifi27l2a), some Cluster 2 microglia (Lpl), or small subpopulations of Cluster 2 microglia (Ccl4 (2a+2b), Cxcl10 (2b), Birc5 (2d).

To capture microglia responding to the injury, white matter was microdissected from LPC- and saline-injected adult (P100) mice and processed using a previously described protocol (Fig 6a). Uninjected P100 whole-brain control samples were collected and processed in parallel. Analysis of microglia from these three conditions produced two major clusters (Fig 6b). Injury Cluster 1 was composed mainly of whole brain and saline-injected control microglia, whereas Injury Cluster 2 was almost entirely composed of microglia from LPC-injected demyelinated lesions (Fig 6c). Microglia in Injury Cluster 2 had downregulated expression of the canonical microglial markers *P2ry12* and *Cx3cr1,* a phenotype that has been observed in other injuries and diseases (Haynes et al., 2006; Keren-Shaul et al., 2017; Salter and Stevens, 2017). Interestingly, we found that Injury Cluster 2-specific genes were variably upregulated among the microglia (Fig 6d), suggesting the existence of subpopulations within the cluster. To delineate these genes, we created three categories: broadly responsive genes that were upregulated in greater than 60% of Injury Cluster 2 microglia, responsive genes that were upregulated in 30-60%, and selectively responsive genes that were upregulated in less than 30% (Fig 6d). Broadly responsive genes included apolipoprotein E (*Apoe), Ifi27l2a*, and the major histocompatibility complex II (*MHC-II*) genes *H2-Aa* and *H2-K1* (Fig 6d). Responsive genes included several interferon response genes (*Irf7, Oasl2*, and *Ifit3), Ccl3,* and lipoprotein lipase (*Lpl*) (Fig 6d). Selectively responsive genes included AXL receptor tyrosine kinase (Axl), *Ccl4*, chemokine (C-X-C motif) ligand 10 (*Cxcl10*), and *Birc5* (Fig 6d).

To further investigate microglial subpopulations within the lesion, we performed a second level of clustering analysis called subclustering on only Injury Cluster 2 cells (Fig 6e) and found several small subpopulations of injury-responsive microglia (Injury Subclusters 2a-d, Fig 6e). Injury Subcluster 2d expressed cell proliferation markers, including *Birc5,* a gene also expressed by proliferative microglia during early brain development (Fig 4g). Microglia from Injury Subcluster 2b were specifically enriched for the interferon response gene *Cxcl10* (Fig 6e). Two of the Injury Subclusters 2a and 2b were specifically enriched for *Ccl4* (Fig 6e). In addition to these distinct transcriptional programs, each Injury Subcluster upregulated the broadly responsive genes, including *Apoe* (Fig 6g). These results suggest that microglial activation is a tailored response in which microglia activate both generalized and selective transcriptional programs.

Demyelinated lesions contain a high density of activated microglia, generally in the core, where they can remove myelin debris and release factors into the lesion microenvironment. Given the variable gene response in the Injury Cluster 2 microglia and the different subpopulations of activated microglia found in our subclustering analysis, we performed smFISH to determine the distribution of these microglia within the injured tissue (Fig 7a-d). A comparison of saline-injected controls and LPC-injected animals showed a large increase in the number of *Fcrls*^+^ microglia throughout the demyelinated tissue, as well as a broad upregulation of *Apoe* in microglia (Fig 7a), which supports our sequencing data (Fig 6d). Surprisingly, smFISH showed that two of the selectively responsive genes - *Ccl4* and *Cxcl10* - were confined to small patches within the larger lesion (Fig 7b). Both markers colocalized with *Fcrls* in many, but not all cases, suggesting other cells might also upregulate these pathways. Very few *Ccl4+Fcrls*^+^ and *Cxcl10+Fcrls*^+^ cells were found in saline-injected controls compared to LPC-injected lesions, suggesting that these effects were in response to demyelination and were not a consequence of the injection (Fig 7c).

**Figure 7.**
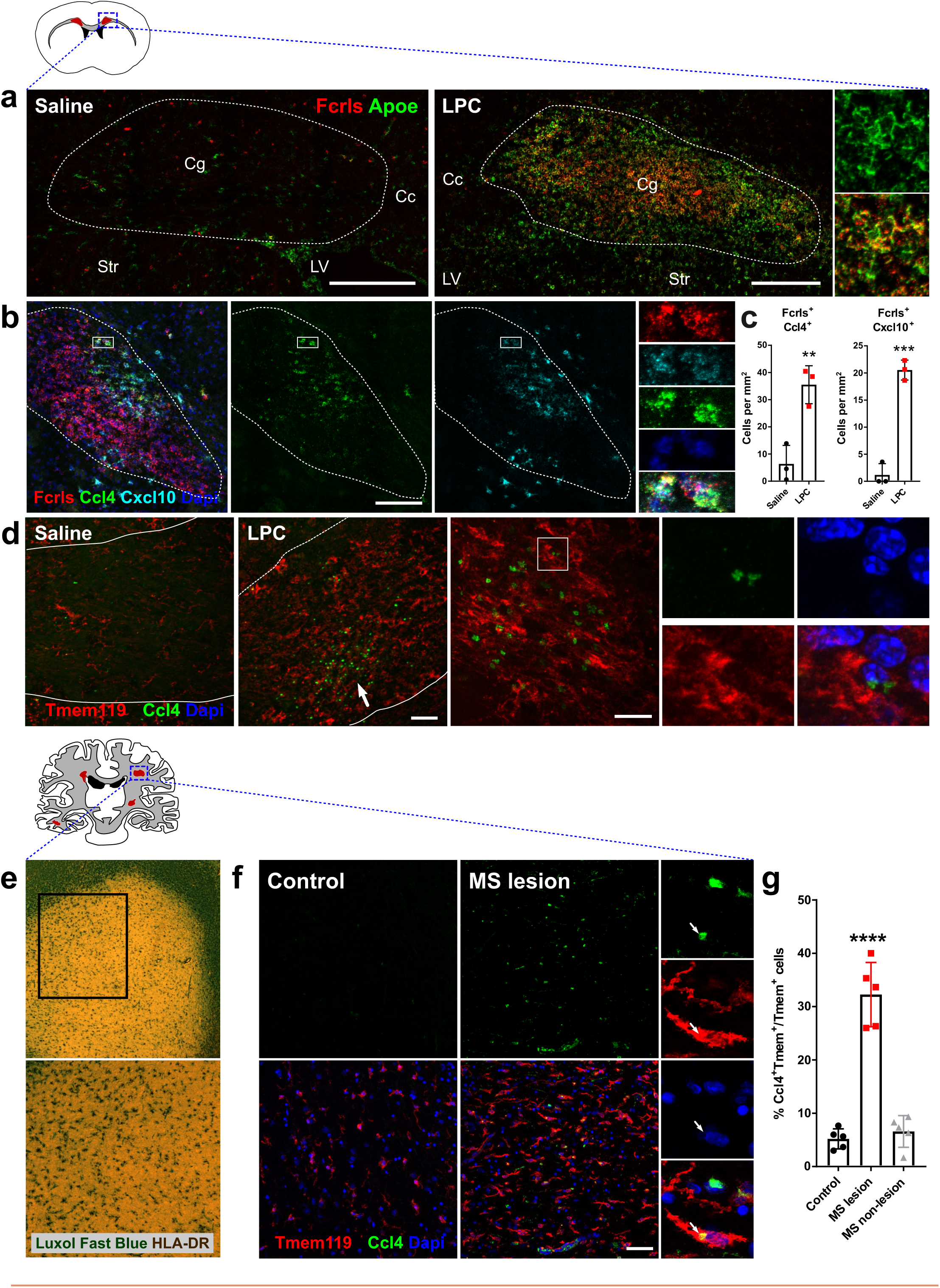
Different subpopulations of activated microglia are found in spatially restricted areas of mouse demyelinated lesions and human MS lesions. (a) Low-magnification confocal images of saline- and LPC-injected white matter in the corpus callosum and cingulum in adult mice stained by smFISH for the microglial probe Fcrls and broadly responsive microglia activation marker Apoe. Scale bar = 250 microns. (b) Confocal image of LPC-injected demyelinated lesion stained by smFISH for the probes Fcrls and selectively responsive microglia activation markers Ccl4 and Cxcl10. Scale bar = 100 microns. (c) Quantification of the percent of Fcrls+ microglia in saline- and LPC-injected white matter that co-expressed the selectively responsive microglia activation markers Ccl4 and Cxcl10. N = 3 mice, ***P<0.001, **P<0.01, unpaired t-test. (d) Confocal images of saline- and LPC-injected white matter stained with antibodies to recognize Ccl4 and resident microglia marker Tmem119. Small scale bar (left) = 50 microns, bigger scale bar (right) = 20 microns. (e) Low-magnification images of demyelinated active MS lesions stained with Luxol Fast Blue (myelin) and anti-HLA-DR antibody (microglia/immune cells). (f) Confocal images of control and human MS white matter stained with antibodies to recognize Ccl4 and resident microglia marker Tmem119 and DAPI+ cell nuclei. Scale bar = 50 microns. (g) Quantification of the percent of Ccl4+Tmem119+ microglia in control patients, active MS lesions, and unaffected white matter in MS patients. N = 5 patients per condition, ****P<0.0001, one-way ANOVA, Tukey’s post-hoc analysis

To identify these subpopulations at the protein level, we stained saline- and LPC-injected lesions with anti-Ccl4 and anti-Tmem119, a marker selective to resident microglia and not downregulated at the protein level following injury (Bennett et al., 2016). We found a large increase in the number of *Tmem119*^+^ microglia in the core of the lesion, as compared to saline-injected controls (Fig 7d). We also found a striking increase in the number of Ccl4+ cells (Fig 7d), which formed patches within the lesions, similarly to the smFISH analysis (Fig 7b). *Ccl4* staining was found in *Tmem119*^+^ cells and was perinuclear, most likely in the Golgi, where cytokines are often packaged into vesicles for release.

*Ccl4* is upregulated in the brains of Multiple Sclerosis (MS) patients (Szczucinski and Losy, 2007), where it could govern both trafficking and effector functions of *CCR5*^+^ immune cells. MS is caused, in part, by immune-cell infiltrates that cause myelin degeneration, however the mechanisms that trigger infiltration and the role of microglia in stimulating this process are not well understood. To test whether we could identify a *Ccl4*^+^ subset of microglia in human MS lesions, we co-stained active demyelinated lesions from five age-matched control and patient samples (Fig 7e). We used a human-specific anti-Tmem119 antibody to mark resident microglia, since its expression is maintained by resident human microglia in MS lesions and is not expressed by infiltrating immune cells (Zrzavy et al., 2017). Co-staining with human anti-Ccl4 antibody showed that Ccl4 was upregulated in the core of active lesions, compared to unaffected white matter in control patients (Fig 7f). The percentage of Ccl4+ cells away from the active lesions was comparable to that seen in control patient tissue, suggesting this specific subset of microglia is only present in the lesion (Fig 7g). Ccl4 staining was found predominantly in Tmem119^+^ microglia, although a small percentage of Ccl4+/Iba1+ amoeboid immune cells were also observed (data not shown). Ccl4+ cells were also found in blood vessels in control and MS samples (as seen at the bottom of the MS lesion image, Fig 7f). In total, approximately 30% of microglia in the lesion were *Ccl4*^+^ (Fig 7g). These results suggest that specific subpopulations of microglia are similarly represented in mouse lesions and MS white matter lesions and that they are the predominant source of *Ccl4* in MS lesions. Detailed analysis of whether the other genes enriched in these cells in mice translate to this subpopulation in humans will confirm their unique identity and inform strategies to manipulate them in MS.

Altogether, these results suggest that microglia ‘activation’ is a dynamic response involving transcriptionally and spatially distinct subpopulations. We were able to translate these findings to human disease, which could mean that markers of these unique - and potentially pathogenic - microglia could be used as biomarkers or novel therapeutic targets. The transcriptional signatures presented here will allow functional interrogation of each responsive microglial state and will help redefine how we classify microglial activation states *in vivo*.

## Discussion

The capacity of microglia to sense and respond to changes in their environment is one of their defining features, but how diverse these responsive states are, when and where they occur in the brain, and how they differ molecularly and functionally has been largely unknown. In our study, we present a comprehensive atlas of microglial heterogeneity that uncovers microglial states that change over development, with age, and in injury. High-throughput droplet-based single-cell sequencing enabled us to analyze the transcriptomes of tens of thousands of cells and to identify small subpopulations of mouse microglia. Our results suggest that microglia assume many distinctive states that change over time, states that can be defined by unique markers and localized within the brain. Because delineating closely related cell states using current computational methods can be difficult and over-clustering is a salient concern for these studies, we validated virtually all of the subpopulations using smFISH and antibody staining, demonstrating that this technology can be used to both find populations and characterize their distribution. Furthermore, our detection of these cells in tissue using a different technique confirmed that the isolation of cells did not cause transcriptional artifacts. Our cell clustering accurately represented the different microglial subpopulations *in vivo*. By taking a conservative approach coupled with detailed histology, we were able to identify and localize novel microglial subpopulations. These newly described transcriptional signatures will provides the foundational information required for allow the development of new tools - including new *Cre* driver lines - to interrogate microglial function and offer a shift in the current paradigm of how we understand, classify, and delineate microglial populations throughout development - as well as in disease, where changes in microglial morphology and ‘activation’ state have been observed but not understood.

### Identification of distinct subpopulations of microglia that suggest functional states in the developing brain

Microglia constantly survey the brain by extending and retracting their processes and assume diverse morphologies that have long been used in histological studies to describe differences in microglial state; for example, in early development, microglia have fewer processes, by adulthood, they are highly ramified, and in disease, these processes can disappear almost completely (Karperien et al., 2013). While these observations undoubtedly hint at underlying biology, the morphology alone tells us very little about their function. Furthermore, a lack of unique markers has further limited our ability to classify these physical transformations. We defined at least 9 subpopulations of microglia and provide unique transcriptional and spatial signatures that give us hints about their possible functions. Microglia development had been previously categorized in bulk RNA-seq studies, but our new cluster definitions highlight the greater degree of developmental complexity that were covered up by dominating signatures in bulk sequencing experiments. These signatures - cell cycle, phagocytosis, and surveillance - correspond with only three of the nine states presented here (Matcovitch-Natan et al., 2016). The specific roles of each microglial state will need to be tested directly, using genetic manipulation and other tools as they become available. This can be done by targeting each group of cells or on a gene-by-gene basis in each subpopulation and will provide a deeper mechanistic insight into microglia signaling mechanisms.

We found that microglial diversity is highest during early development, when microglia are still differentiating (Matcovitch-Natan et al., 2016). Recent evidence has shed light onto some of the signals, including *TGF-beta,* that direct this differentiation process, but the speed at which microglia mature and the number of states they assume during this time was not previously known. Here, we identified a unique microglial state (Cluster 6) that was enriched at E14.5 and defined by the genes *Ms4a7*, *Ccr1, and Mrc1*, which are also enriched in brain border macrophages. These microglia, which were present in the brain parenchyma, could be microglia that have yet to fully differentiate. In support of this idea, blocking *TGF-beta* signaling (which helps to confer microglia identity, (Butovsky et al., 2014)) has no major effect on microglia cell number, but causes a drastic shift in their gene expression including widespread upregulation of (or failure to downregulate) genes found in Cluster 6 microglia and in brain border macrophages, including *Mrc1* (Wong et al., 2017). These microglia could also arise from ontologically distinct progenitors - recent transplantation experiments suggest that *Ms4a7* and other markers in Cluster 6 are specific to hematopoietic derived cells versus those derived from yolk-sac progenitors (Bennett et al., 2018). Lineage tracing will be essential to understanding their origin and whether they give rise to microglia that persist in the brain. Also, whether Cluster 6 microglia also have specialized functions in brain development remains to be seen, but their enrichment for several members of the Ms4a gene family, which are risk genes for Alzheimer’s disease (Ma et al., 2015), could confer unique functional properties and necessitates further exploration.

Other key microglial states in early development were enriched for pathways associated with cell metabolism, growth, and motility (Cluster 3) and proliferation (Clusters 2a-c) and were scattered throughout the brain. Clusters 2 and 3 had considerable transcriptional overlap, suggesting that microglia transition in and out of both states. Because microglia migrate, grow, and proliferate in response to injury, teasing apart the molecular triggers regulating these pathways could allow for the manipulation of microglial number as a treatment - for example, to reduce the number of reactive microglia and limit inflammation, astrogliosis, and cytotoxicity in neurodegenerative disease (Liddelow et al., 2017). Furthermore, these cells had a distinct metabolic signature. Metabolic state is directly manipulated by cytokine and pattern-recognition receptors, and elevated glycolytic activity has been tied to immune cell activation (Everts et al., 2014; O’Neill and Pearce, 2016; Pearce and Everts, 2015). Processing fatty acids also plays an integral role in controlling the macrophage activation state, a process that could be mediated by *Fabp5* in microglia (Jha et al., 2015). However, it remains to be seen whether metabolic activity of Cluster 3 microglia is most reflective of their growth, proliferation, motility, or activation state.

The other microglial subpopulation in the developing brain, Cluster 4, was restricted to unmyelinated axon tracts at P4/P5 and assumed a highly activated profile, which was surprising in the absence of any pathology. The temporally and spatially restricted nature of these cells suggests extrinsic forces shape this state, but what those signals are and where they come from is still unknown. This subpopulation of activated microglia was most likely first described in 1932 by Pio del Rio-Hortega and dubbed the “fountains of microglia” due to their density and amoeboid morphology around the brain ventricles - as if they were streaming into the brain. Two recent studies have linked these cells to myelin formation and hypothesized that they release myelinogenic molecules, but this hypothesis has not been directly tested by selectively targeting these cells (Hagemeyer et al., 2017; Wlodarczyk et al., 2017). Another group showed that these microglia were the predominant source of *Igf1* in the early postnatal brain, and *Igf1* conditional knockout in microglia affected the health of developing cortical neurons at the same age (Ueno et al., 2013). These studies point to an important developmental role for these cells, which can be further explored using the highly defined transcriptional signature we present here.

### Localization of distinct subpopulations of microglia in aging and injured brain

Pathways expressed in development are often reexpressed in disease, and it has long been hypothesized based on morphological similarities between developing microglia and activated microglia that injury causes a reversion to an immature state. We found that microglia responding to injury do re-express developmental markers, but their transcriptional states do not completely overlap. The greatest similarity in cell state was found between P5 Cluster 6 microglia and microglia in the demyelinated lesion. Both groups of cells upregulated similar genes including *Apoe, Lpl,* and *Spp1.* These markers may comprise elements of a common activation pathway, since these genes are also upregulated in microglia that surround plaques in an Alzheimer’s disease model (Keren-Shaul et al., 2017). That microglia are activated by comparable signaling mechanisms in development (Cluster 6) and disease/injury is a tantalizing and widely discussed concept. Our characterization of their transcriptomes will enable investigation of the triggers that cause this response in both contexts. In addition to this common activation state, we also found that injury-responsive microglia upregulated other pathways found in developmental subpopulations including markers of proliferation (*Birc5*), metabolic activity (Cluster 3 microglia *[Mif]*), and those belonging to the Ms4a family (*Ms4a6c).* Re-expression of these markers could be an integral part of the activation process and might be necessary to stimulate a quick response to pathology. Therefore, as we learn more about the role of these pathways in development, we will be able to assess their importance in disease.

A major finding in our study was a novel subset of microglia that selectively express the chemokine *Ccl4.* These microglia are present at low levels during development and expand in two different contexts: aging and injury. The role of inflammatory molecules in the brain has long been studied, and it has been assumed, based largely on *in vitro* studies, that microglia are a major source of these factors (Ransohoff, 2016). However, we found that the only microglia enriched for inflammatory signals are in Cluster 8/Aging Cluster 2/Injury Subcluster 2a,b and express *Ccl3, Ccl2, Ccl7, Ccl9, Ccl12, Il1b* and *Tnf.* Since this small subpopulation of microglia was present at low levels throughout the mouse lifespan, it is possible that they are a specialized group uniquely primed to produce an inflammatory response. Many of the signals expressed in this subpopulation can be highly damaging to the brain. For example, *Il1b* and *Tnf* can both cause neurotoxicity (Takeuchi et al., 2006; Ye et al., 2013), and infiltrating immune cells attracted by the chemokines expressed in the *Ccl4*^+^ subpopulation can exacerbate pathology (Gadani et al., 2015). In MS, infiltrating immune cells specifically target myelin and cause white matter lesions. Precise targeting of this inflammatory subpopulation of microglia in disease could yield a safe and effective way to limit the negative side effects of microglial activation while maintaining the beneficial functions including removal of myelin debris and release of neurotrophic signals (Miron et al., 2013).

Microglia are an especially attractive target for biomarker development and to monitor clinical progression because they can sensitively respond to changes in the brain that emerge before physical symptoms. For example, microglia become activated and aberrantly remove synapses in the early stages of pathology in mouse models of Alzheimer’s disease, well before cognitive decline and plaque formation (Hong et al., 2016). In fact, evidence of aberrant microglia activation has been found in many forms of neurodevelopmental and neurodegenerative disease, but the diversity of these responses and specificity of the microglia response to each disease needs further exploration (Salter and Stevens, 2017). Current PET imaging ligands, including TSPO, are not selective for microglia and are better markers of neuroinflammation (Pappata et al., 1991; Politis et al., 2012). Therefore, identification of robust markers for different microglia states in disease will allow for the development of tools to more specifically track and visualize microglial activation, inflammation, and disease progression.

## Contributions

T.R.H carried out most of the experiments including both the wet lab experiments and bioinformatics analysis, with help from co-authors. C.D. performed all the smFISH and developed the automated smFISH analysis pipeline. L.D-O. and A.J.W. performed flow cytometry, helped to conceptualize experiments and generated figure illustrations. S.G. performed the LPC lesion surgeries. A.Y. and M.S. did the staining and analysis of human MS tissue. F.G. helped with the analysis of the aged brains. A.W., J.N., A.S., and E.M. helped with the single cell sequencing analysis and provided analysis expertise. R.J.M.F helped to design human MS tissue experiments. X.P. helped to design LPC lesion experiments. T.R.H., S.M. and B.S. designed the study and wrote the manuscript.

## Acknowledgements

We would like to thank Dr. Richard Ransohoff, Dr. Christina Usher, Dr. Sam Marsh, and Dr. Daisy Robinton for their critical feedback and insight while preparing the manuscript. We would like to thank Jon Hammond for developing the website and searchable dataset (www.microgliasinglecell.com). This work was funded by grants from the Simons Foundation/SFARI (#346197 to S.M. and B.S.), Rettsyndrome.org mentored fellowship training award (#3214 to T.H.), Silvio O. Conte Center (NIH #P50MH112491 to B.S. and S.M.), NMSS Postdoctoral fellowship Award FG 2063-A1/2 (S.G.), Stanley Center for Psychiatric Illness at the Broad Institute (B.S. and S.M.), and Helen Hay Whitney Fellowship (A.S.).

